# Dilated cardiomyopathy variant R14del increases phospholamban pentamer stability, blunting dynamic regulation of cardiac calcium handling

**DOI:** 10.1101/2023.05.26.542463

**Authors:** Sean R. Cleary, Allen C. T. Teng, Audrey Deyawe Kongmeneck, Xuan Fang, Taylor A. Phillips, Ellen E. Cho, Peter Kekenes-Huskey, Anthony O. Gramolini, Seth L. Robia

## Abstract

The sarco(endo)plasmic reticulum Ca^2+^ ATPase (SERCA) is a membrane transporter that creates and maintains intracellular Ca^2+^ stores. In the heart, SERCA is regulated by an inhibitory interaction with the monomeric form of the transmembrane micropeptide phospholamban (PLB). PLB also forms avid homo-pentamers, and dynamic exchange of PLB between pentamers and the regulatory complex with SERCA is an important determinant of cardiac responsiveness to exercise. Here, we investigated two naturally occurring pathogenic mutations of PLB, a cysteine substitution of arginine 9 (R9C) and an in-frame deletion of arginine 14 (R14del). Both mutations are associated with dilated cardiomyopathy. We previously showed that the R9C mutation causes disulfide crosslinking and hyperstabilization of pentamers. While the pathogenic mechanism of R14del is unclear, we hypothesized that this mutation may also alter PLB homo-oligomerization and disrupt the PLB-SERCA regulatory interaction. SDS-PAGE revealed a significantly increased pentamer:monomer ratio for R14del-PLB when compared to WT-PLB. In addition, we quantified homo-oligomerization and SERCA-binding in live cells using fluorescence resonance energy transfer (FRET) microscopy. R14del-PLB showed an increased affinity for homo-oligomerization and decreased binding affinity for SERCA compared to WT, suggesting that, like R9C, the R14del mutation stabilizes PLB in its pentameric form, decreasing its ability to regulate SERCA. Moreover, the R14del mutation reduces the rate of PLB unbinding from the pentamer after a transient Ca^2+^ elevation, limiting the rate of re-binding to SERCA. A computational model predicted that hyperstabilization of PLB pentamers by R14del impairs the ability of cardiac Ca^2+^ handling to respond to changing heart rates between rest and exercise. We postulate that impaired responsiveness to physiological stress contributes to arrhythmogenesis in human carriers of the R14del mutation.

## INTRODUCTION

Dilated cardiomyopathy (DCM) is the most common disorder leading to heart failure, characterized by thinning of the ventricular wall and reduced pumping capacity of the heart (*1, 2*). As much as 50% of DCM is thought to be caused by genetically inherited mutations (*1, 2*). Although DCM mutations predominantly occur in proteins of the sarcomere, cytoskeleton, and nuclear membrane (*1, 3, 4*), a number of mutations have been identified in phospholamban (PLB), a transmembrane micropeptide that regulates intracellular Ca^2+^ transport by the sarcoplasmic reticulum (SR) Ca^2+^ ATPase (SERCA) (*5-9*). SERCA sequesters cytosolic Ca^2+^ into the SR lumen to relax cardiac muscle between beats and prepares an intracellular Ca^2+^ store that is released into the cytoplasm to coordinate subsequent contractions of the heart (*10*). Dysregulation of SERCA Ca^2+^ transport is thought to cause maladaptive remodeling of intracellular Ca^2+^ signaling pathways that eventually lead to heart failure in patients (*11, 12*).

PLB is an SR-localized transmembrane peptide consisting of 52 amino acids that form 2 α-helical domains: a transmembrane helix that spans a single pass through the SR membrane and a cytoplasmic domain (*13*). PLB inhibits SERCA function through a direct protein-protein interaction between the transmembrane domain of PLB and a regulatory cleft within SERCA’s transmembrane domain (*14, 15*). The bioavailability of free PLB monomers that can regulate SERCA is dynamically controlled by PLB homo-oligomeric assembly into PLB pentamers (*16-19*). Naturally occurring pathogenic mutations in the cytoplasmic domain of PLB have been shown to alter the stability of PLB pentamers, causing chronic dysregulation of SERCA leading to DCM (*20-22*). In particular, we previously showed an autosomal-dominant mutation of arginine 9 to a cysteine (R9C) causes disulfide bonds to form under conditions of oxidative stress (*22*). This crosslinking stabilizes PLB in pentamers, chronically reducing the population of active monomeric PLB available to regulate SERCA (*22*). Conversely, another autosomal dominant DCM mutation, which causes an in-frame deletion of arginine 14 (R14del), has been proposed to have the opposite effect. R14del-mutant PLB monomers were reported to destabilize PLB pentamers into smaller oligomer species when heterologously expressed with WT-PLB, with consequent super-inhibition of SERCA (*5*). However, several subsequent studies showed that the R14del mutation decreases the inhibitory potency of PLB (*21, 23-25*), so the pathogenic mechanism of this mutation remains unclear.

Aside from a steady-state effect on the PLB monomer-pentamer equilibrium, DCM-associated mutations may also impact the heart’s dynamic responsiveness to physiological stress such as exercise. Phosphorylation of PLB by PKA during adrenergic stimulation of the heart relieves PLB inhibition, in part through stabilization of PLB pentamers (*16, 17, 19*), providing a mechanism to temporarily increase SERCA Ca^2+^ transport during exercise or other physiological stress. However, both R9C and R14del display reduced responsiveness to phosphorylation (*5, 20, 22, 26, 27*). In addition, we showed that temporal buffering of the concentration of PLB monomers by the pentamer controls the responsiveness of PLB regulation to changing heart rates (*19*). Specifically, we showed that a fraction of PLB unbinds from SERCA in response to cellular Ca^2+^ elevations during cardiac systole (contraction), and this dynamic fraction of PLB is rapidly incorporated into PLB pentamers. This process reverses itself during diastole (cardiac relaxation) as Ca^2+^ levels fall, but we (*18, 19*) and others (*28*) showed that the rate of PLB pentamer dissociation is slow. This limits the rate of return of PLB monomers to SERCA during diastole. Because of this “kinetic trapping” mechanism, PLB increasingly accumulates in pentamers as the heart rate increases, which helps to relieve inhibition of SERCA during exercise. This mechanism may contribute to the Bowditch effect, the positive relationship between cardiac force and pacing frequency (*19*). Since both R9C and R14del mutations have been proposed to alter the stability of PLB pentamers, it seems likely that both the equilibrium regulation of SERCA and the ability of PLB regulation to adjust to changing physiological demands may be impacted by these mutations. Here, we used a combination of biochemical and biophysical measurements to study the impact of R9C and R14del on PLB binding equilibria. The results suggest both mutations may dysregulate SERCA function through similar pathogenic mechanisms.

## RESULTS

### R14del-PLB forms more stable pentamers than WT-PLB

We previously reported that a naturally-occurring mutation of PLB linked to DCM, R9C, stabilizes PLB pentamer interactions through oxidative cross-linking of mutant cysteines (*22*). It, however, remains unknown if the R14del mutation, which is also associated with DCM, also alters PLB oligomerization. To test this hypothesis, we first monitored the oligomerization status of WT-PLB, WT-PLB : R14del-PLB (1:1), and R14del-PLB complexes by immunoblots. **Fig. 1A** showed the expected mix of monomers and oligomers of WT-PLB in homogenates. Unexpectedly, we observed increased oligomerization for R14del-PLB compared to WT-PLB. Mixed hetero-oligomers of R14del- and WT-PLB also showed increased high molecular weight bands compared to WT-PLB homo-oligomers. Next, we tested the thermal stability of the oligomeric species subjecting WT- and R14del-PLB oligomers to increased temperature. Immunoblotting revealed increased stability of R14del-PLB compared to WT-PLB across the tested range of temperature (**Fig. 1B**). Quantification of the oligomer bands revealed 87.2 ± 3.2% oligomers for R14del-PLB versus 79.5 ± 0.8% for WT-PLB at 25 °C, and the R14del-PLB melting curve was right-shifted compared to WT-PLB (**Fig. 1C**). These surprising findings suggested that R14del-PLB is more oligomeric than WT-PLB.

**Figure 1.**
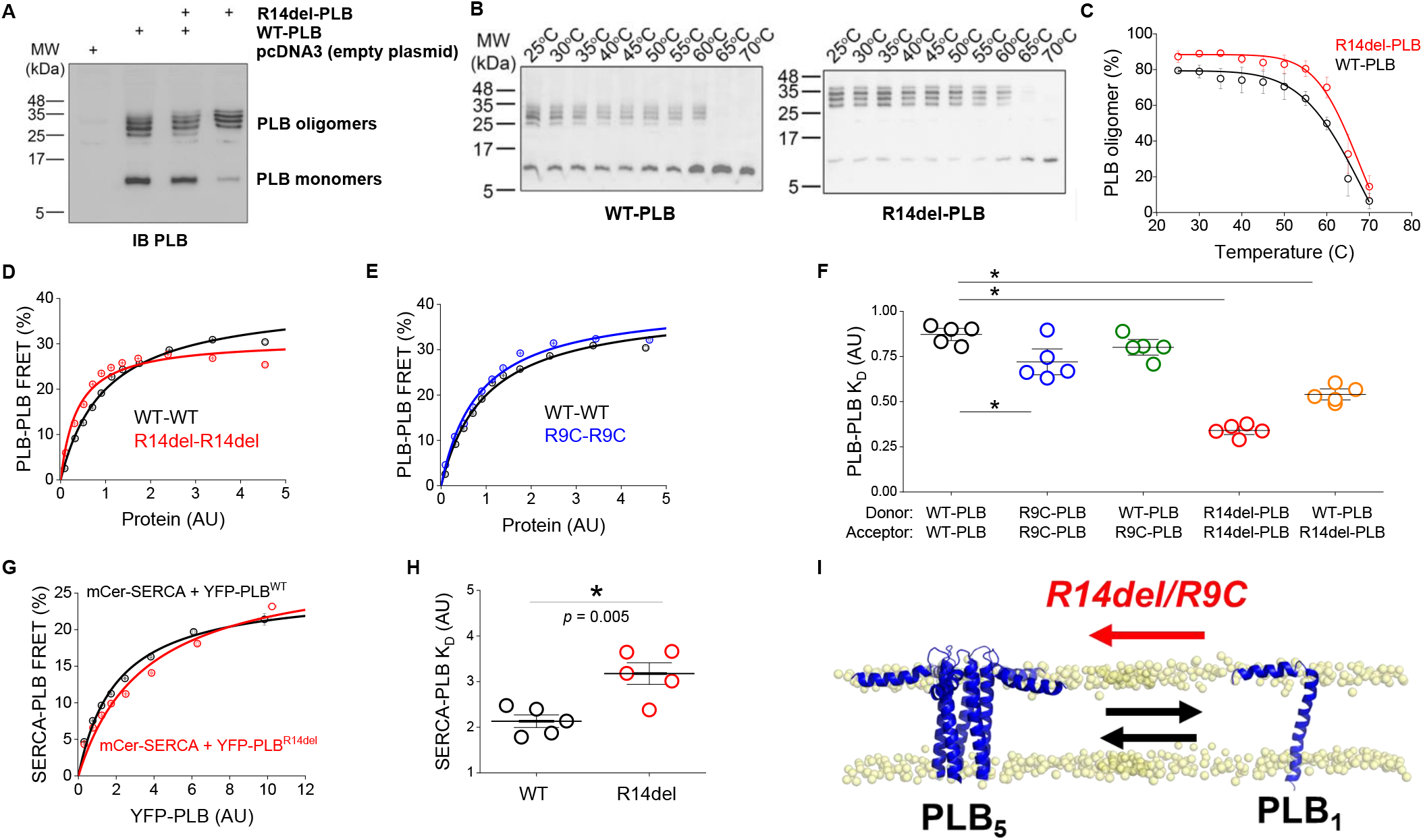
R14del-PLB mutation stabilizes PLB pentamers. **A)** An immunoblot showing increased oligomers (25 kDa and above) and reduced PLB monomers (7 kDa) for R14del-PLB vs. WT-PLB. **B)** PLB thermostability assays showing that PLB pentamers (WT-PLB, left panel; R14del-PLB, right panel) dissociated into monomers with increasing temperature (25-70 °C). **C)** Quantification of immunoblots yielded oligomer melting curves. R14del-PLB oligomers (*red*) were more stable than PLB-WT oligomers (*black*) between 25 and 70 °C (*n* = 3). **D)** FRET-based binding curves reveal a left shift in the concentration dependence of the PLB-PLB interaction with R14del mutation. **E)** FRET-based binding curves reveal a left shift in the concentration dependence of the PLB-PLB interaction with R9C mutation. **F)** Apparent K_D_s of PLB pentamer interactions with WT- and R14del-PLB. Differences determined by 1-way ANOVA with Tukey’s post-hoc (* = *p*<0.05). See Supplementary **Table S1** for complete statistical analysis. **G)** A right shift in the concentration dependence of R14del-PLB-SERCA FRET suggested decreased binding of R14del-PLB to SERCA. **H)** Apparent K_D_s of SERCA interactions with WT- and R14del-PLB. Differences determined by student’s t-test (* = *p*<0.05). **I)** A schematic diagram showing that both R14del and R9C mutations both impact the monomer/pentamer equilibrium by shifting the equilibrium towards the PLB pentamer. PDBs: PLB_5_ - 2KYV; PLB_1_ – 1FJP.

To compare the concentration-dependent binding affinity of R14del and WT pentamer interactions in membranes of living cells, we quantified fluorescence resonance energy transfer (FRET) between donor (mCer)- and acceptor (YFP)-labeled PLB expressed in HEK-293 cells. Specifically, using high throughput cell scoring of the fluorescence intensity of YFP-PLB (as an index of protein expression) and quantitative FRET efficiency in transfected cells, we measured the concentration-dependence of PLB pentamer binding for the WT and R14del PLB. Both WT and R14del pentamer FRET increased with increasing protein concentration until the interactions saturated at a maximum FRET value (FRET_max_) (**Fig. 1D**). The data were well described by a hyperbolic fit of the form FRET = (FRET_max_ x [YFP-PLB])/(K_D_ + [YFP-PLB]), where FRET_max_ is the intrinsic FRET of the bound pentamer at high protein concentration and the K_D_ is the apparent dissociation constant, which is inversely related to the affinity of the PLB pentamer. The FRET_max_ for PLB pentamers was consistently decreased for the R14del-R14del interaction compared to WT (**Fig. 1D**), suggesting a small change in the structure of the pentamer. The K_D_ of the R14del-R14del interaction was significantly lower than that of WT-WT (*p* = 1.479 × 10^−8^), consistent with SDS-PAGE and thermal denaturation measurements that suggested R14del oligomerizes with higher affinity than WT pentamers (**Fig. 1A, B**). We also quantified the affinity of another DCM mutation, R9C, which we previously showed increases pentamer binding affinity (*22*). Consistent with that previous study, we found R9C mutation increases the FRET_max_ while significantly reducing the K_D_ the PLB-PLB interaction (*p* = 0.012) (**Fig. 1E,F**), indicating a change in the R9C pentamer structure and an increase in oligomerization affinity, respectively. Since patients that carry R14del and R9C mutations are heterozygous, expressing both WT and mutant copies of the PLB gene, we also quantified FRET between WT-PLB labeled with a fluorescent donor and mutant PLB labeled with a fluorescent acceptor. Interestingly, the K_D_ for WT-R14del binding was significantly lower than that of the WT-WT interaction (*p* = 0.04) (**Fig. 1F**), indicating that R14del-PLB also binds WT-PLB with higher affinity. The value of the WT-R9C K_D_ was also lower than that of WT-WT, which would be consistent with our previous study (*22*), though the difference did not achieve statistical significance here. Taken together, the results suggest that both DCM mutations of PLB oligomerize more avidly and this is a dominant effect, increasing sequestration of WT-PLB protomers together with mutant protomers into avidly bound, mixed pentamers.

We also measured the effects of R14del mutation on the PLB-SERCA interaction by measuring FRET between donor labeled SERCA and acceptor labeled WT- and R14del-PLB (**Fig. 1G**). The FRET_max_ of SERCA binding with WT-PLB was significantly lower than the interaction of the Ca^2+^ pump with R14del-PLB (*p* = 0.01) (**Fig. S1**), suggesting a shorter distance between the fluorescent tags fused to the N-termini of R14del-PLB and SERCA. The K_D_ of SERCA-PLB binding was significantly higher with R14del-PLB than WT (*p* = 0.005) (**Fig. 1G**), indicating decreased binding to SERCA. This is probably not due to a true change in the intrinsic affinity of the R14del-PLB monomer for SERCA, rather, we attribute it to decreased bioavailability of the mutant due to increased sequestration into pentamers. The results from fitting FRET-based binding curve data are summarized in **Table 1**. Overall, the data indicate that both R9C and R14del mutations cause PLB to form more stable pentamers at the expense of binding to SERCA.

**Table 1.** PLB regulatory interactions quantified by FRET.

| Binding Partners<br>(Donor – Acceptor) | FRET<sub>max</sub> (%) | Apparent K<sub>D</sub> (AU) |
| --- | --- | --- |
| WT-PLB – WT-PLB | 39.0* | 0.87 ± 0.05 |
| R9C-PLB – R9C-PLB | 39.4* | 0.72 ± 0.11 |
| WT-PLB – R9C-PLB | 37.5* | 0.80 ± 0.06 |
| R14del-PLB – R14del-PLB | 31.7* | 0.34 ± 0.03 |
| WT-PLB – R14del-PLB | 34.8* | 0.54 ± 0.05 |
| SERCA – WT-PLB | 24.7 ± 1.0 | 2.13 ± 0.31 |
| SERCA – R14del-PLB | 28.5 ± 2.4 | 3.12 ± 0.53 |

Data are reported as mean ± SD ( $n = 5$ ). \* PLB-PLB FRET-based binding curve data were globally fit to share a single, best FRET<sub>max</sub> value within each group.

Another group previously used solution and solid-state NMR to determine that R14del mutation increased the conformational dynamics of the cytoplasmic domain of PLB. This caused the domain to shift towards an unfolded conformation that was more detached from the membrane. These observations positively correlated with other PLB mutations that cause loss-of-inhibition (*29*). However, that study used a variant of PLB containing additional mutations that destabilized the PLB pentamer, and so the experiments could not reveal structural determinants of increased R14del-PLB oligomer stability. To investigate how R14del mutation might impact the structure or dynamics of PLB pentamers, we generated a model of the R14del-PLB pentamer based on the NMR structure of the WT-PLB pentamer (PDB: 2KYV) (*13*) (**Fig. 2A**), removing Arg14 and bridging the ends of Arg13 and Ala15 while maintaining the value of the torsion angle omega for the 14-15 peptide bond (**Fig. S2**). Inspection of the solvent accessible electric potential of the R14del pentamer showed the expected diminution of positive charge (**Fig. 2B**, *blue*) compared to the WT pentamer structure. The WT pentamer structure and R14del pentamer model were subjected to molecular dynamics (MD) simulations. Deletion of Arg14 initially rotated the helix of the cytoplasmic domain of PLB (amino acids 1-20) by a fraction of a helical turn, but this orientation quickly relaxed at the start of the simulation, and the solvent-exposed hydrophilic residues in the WT structure (including phosphorylation sites S16 and T17) remained solvent-exposed in the R14del model. The relaxation of the cytoplasmic domain helix rotation was facilitated by the highly flexible loop that connects the cytoplasmic domain to the top of the transmembrane helix (amino acids 21-52) (**Fig. S2D**,**E**). Another group observed increased alpha helicity in this flexible loop region after R14del mutation (*30*). However, our MD simulations did not recapitulate this, and we did not detect differences in alpha helical content of R14del compared to the WT pentamer. Ensembles of representative snapshots (**Fig. 2C**) showed that the pentameric arrangements of protomers in the WT structure (*black*) and R14del model (*red*) were intact on the timescale of these simulations. **Fig. 2D** shows the most frequently sampled structures for WT (*black*) and R14del (*red*) pentamers. As expected, the cytoplasmic domain was highly dynamic and the transmembrane region was well-ordered. Indeed, the large amplitude (>5Å) motions of the cytoplasmic domain, which includes the Arg14 locus, made it challenging to detect any differences between the WT and R14del pentamers. Overall, we did not observe clear differences in the 2°, 3°, or 4° structure, or in how the structures evolved over the 1 μs time course of the simulation (**Fig. 2E**), or in structural dynamics as measured from root mean square fluctuation (RMSF) (**Fig. 2F**). We did not detect significant differences in the number of contacts between adjacent protomers in the pentamer (**Fig. S3**) or in the tilt angle of the bundle of transmembrane helices (**Fig. S4A-C**), or in the autocorrelation of time dependent changes in this tilt angle (**Fig. S4D-F**) for R14del compared to WT. Thus, aside from the expected change in electrostatic potential (**Fig. 2B**), possible determinants of the observed increase in R14del-PLB pentamer stability (**Fig. 1**) were not revealed by these μs-timescale simulations.

**Figure 2.**
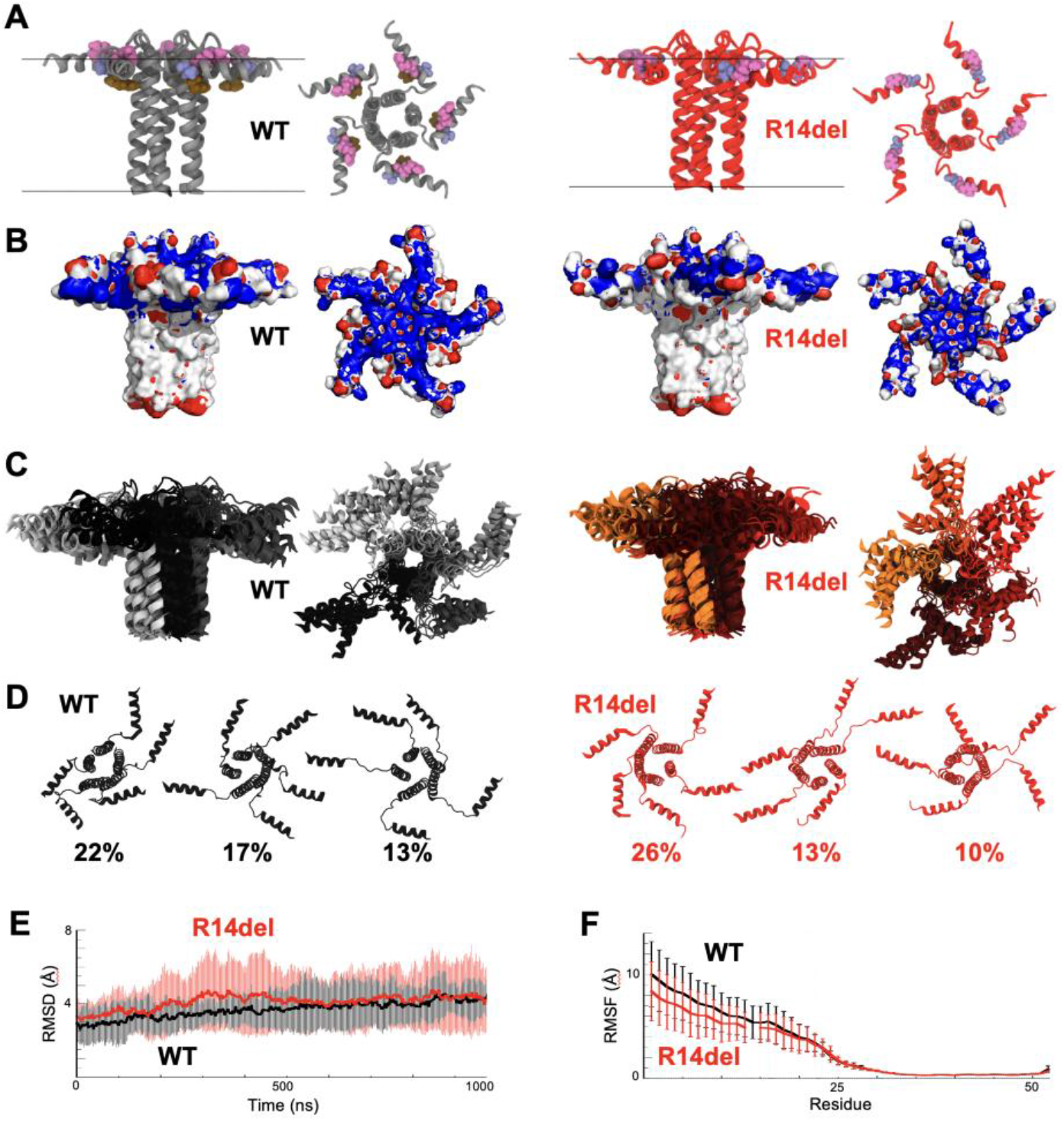
Molecular simulations of PLB pentamer structures. **A)** From left to right: Structure of WT-PLB pentamer (grey, pdb: 2KYV) viewed along the plane of the bilayer and along membrane normal from the cytoplasmic side, compared with a model of the R14del pentamer (red) viewed along the plane of the bilayer and along membrane normal. Membrane surfaces are marked with horizontal lines; space-filling representations highlight residues R13 (pink), R14 (brown) and A15 (blue). **B)** Comparison of the electrostatic potential of WT and R14del pentamers. Positive (blue) and negative (red) isosurfaces represent ± 5KT. R14del showed decreased positive charge in the cytoplasmic domains of the pentamer. **C)** An overlay of representative structures of the WT and R14del pentamers generated from MD simulations. **D)** Classes of structures sampled during MD simulations for WT (black) and R14del (red) determined by cluster analysis. Overall, we did not observe substantial differences in the structure or dynamics of the WT and R14del pentamers. **E)** Pentamer structural evolution during the simulation quantified from RMSD. **E)** Pentamer structural dynamics quantified from RMSF.

### R14del-PLB exchanges slowly from PLB pentamers

We have previously shown that PLB-SERCA complexes are dynamically responsive to cellular Ca^2+^ signals, such that a dynamic fraction of PLB monomers unbind SERCA and rapidly incorporate into PLB pentamers during intracellular Ca^2+^ elevations. In our previous study, we found the rates at which PLB exchanges from pentamers are important for determining the availability of free PLB monomers to regulate SERCA during changing heart rates (*19*). Here, we performed similar measurements to test how stabilization of the pentamer by R9C and R14del mutations may affect the dynamics of PLB exchange from pentamers to SERCA during Ca^2+^ signaling. Briefly, to recapitulate aspects of muscle cell Ca^2+^ signaling in our experimental system, we transfected HEK-293 cells with RyR2 and SERCA2a. These transfected proteins mediate spontaneous ER Ca^2+^ uptake and release events that mimic the dynamic Ca^2+^ signals in cardiac muscle (*19, 31, 32*). Cellular Ca^2+^ elevations were detected by confocal microscopy as an increase in fluorescence of the cytosolic Ca^2+^ indicator dye, X-Rhod-1 (**Fig. 3A,B**, *black*). To monitor dynamic shifts in PLB oligomerization in response to cellular Ca^2+^ elevations, we measured changes in FRET ratio (acceptor/donor) between donor- and acceptor-labeled PLB over time in response to Ca^2+^. Consistent with our previous study, a dynamic fraction of PLB was freed from SERCA and rapidly assembled into pentamers during intracellular Ca^2+^ elevations. We observed this in some cells as an increase in PLB-PLB FRET in response to Ca^2+^ elevations (**Fig. 3A**, *blue*).

**Figure 3.**
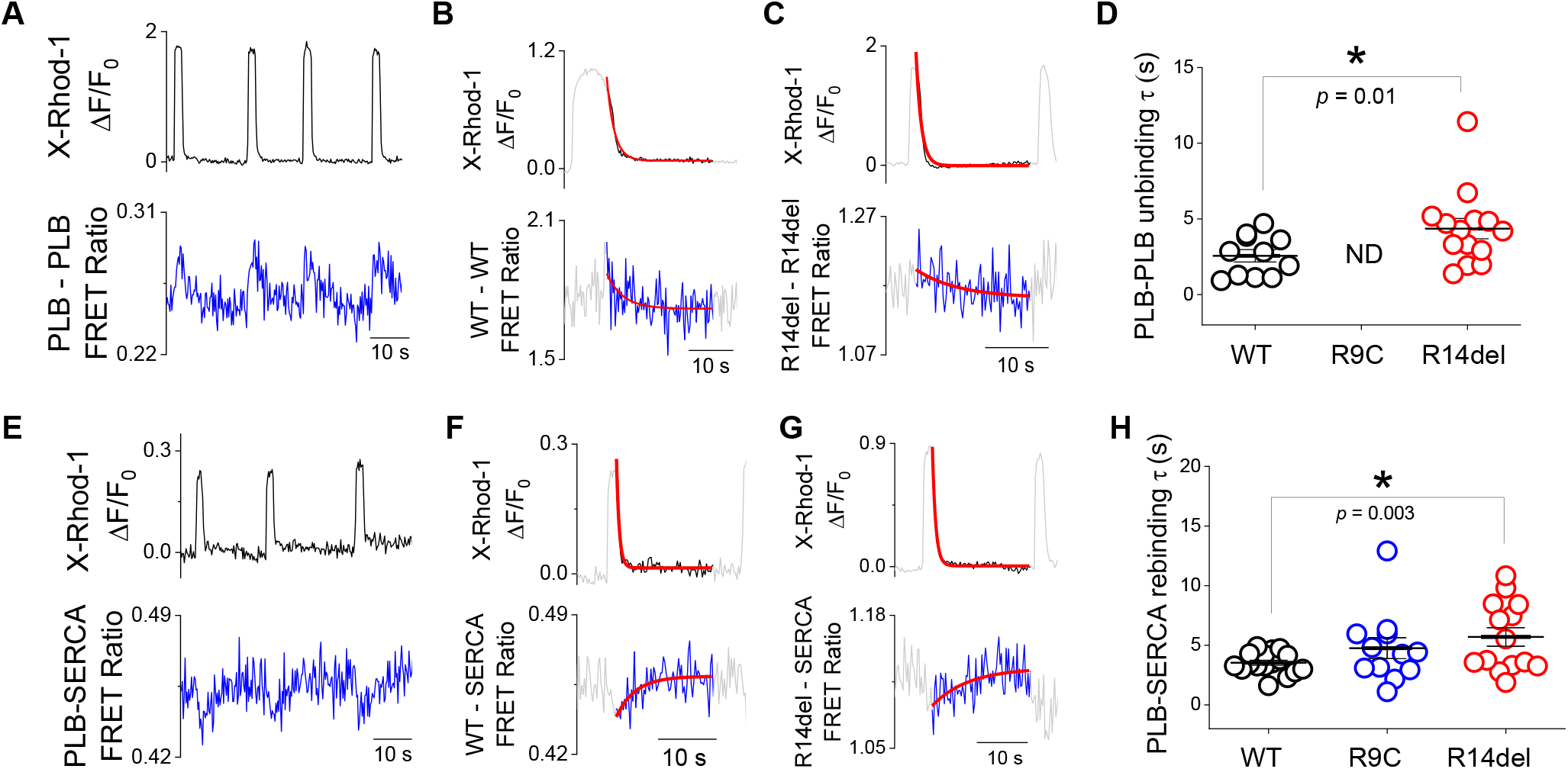
R14del-PLB exchanges slowly from pentamers compared to WT-PLB. **A)** Quantification of X-Rhod-1 fluorescence (*black*) with simultaneous measurement of changes in PLB-PLB FRET (YFP/Cer ratio) (*blue*). **B)** Representative exponential decay fit of human PLB_WT_-PLB_WT_ unbinding during Ca^2+^ uptake. **C)** Representative exponential decay fit of human PLB_R14del_-PLB_R14del_ unbinding during Ca^2+^ uptake. **D)** PLB-PLB unbinding times of WT and mutant pentamers quantified by single-exponential decay fitting. Data are presented as mean ± SE. Differences determined by one-way ANOVA with Dunn’s post-hoc test (* = *p*<0.05). ND = not detected. See Supplementary **Table S3** for complete statistical analysis. **E)** Quantification of X-Rhod-1 fluorescence (*black*) with simultaneous measurement of changes in PLB-SERCA FRET (YFP/Cer ratio) (*blue*). **F)** Representative exponential decay fit of PLB_WT_-SERCA rebinding during Ca^2+^ uptake. **G)** Representative exponential decay fit of PLB_R14del_-SERCA rebinding during Ca^2+^ uptake. **H)** Rebinding times of WT, R9C, and R14del to SERCA quantified by single-exponential decay fitting. Data are presented as mean ± SE. Differences determined by one-way ANOVA with Dunn’s post-hoc test (* = *p*<0.05). See Supplementary **Table S5** for complete statistical analysis.

The rate of PLB oligomerization into pentamers during Ca^2+^ release was rapid, closely matching the rate of the increasing Ca^2+^ signal (**Fig. 3A**). When Ca^2+^ levels decreased to baseline, the unbinding of WT-PLB pentamers lagged behind Ca^2+^ reuptake. Fitting the declining Ca^2+^ and FRET signals to a single exponential function revealed characteristic time constants (τ) for each process. Consistent with our previous study, the τ of WT PLB-PLB unbinding was 2.6 ± 0.4 s, lagging significantly behind Ca^2+^ uptake (τ = 1.0 ± 0.1 s, *p* = 0.01) (**Fig. 3B**). Interestingly, the rate of pentamer unbinding with R14del was significantly slower than WT (τ = 4.4 ± 0.7 s, *p* = 0.01) (**Fig. 3C,D**), suggesting that the increased PLB-PLB interaction stability with R14del mutation causes PLB pentamers to dissociate at a much slower rate. We were not able to detect FRET changes consistent with association/dissociation of the R9C-PLB pentamers (**Fig. 3D**, *ND*). We attribute this to a limited capacity for exchange of PLB from covalently crosslinked pentamers. The kinetics of WT and mutant pentamer unbinding after Ca^2+^ elevations are summarized in Supplementary **Fig. S5** and **Table S2**.

To assess the impact of slower pentamer unbinding on the rates of PLB exchange with SERCA, we measured FRET between donor labeled SERCA and acceptor labeled PLB, comparing the dynamics of WT and mutant PLB acceptors. Consistent with our previous study, we observed dynamic decreases in PLB-SERCA FRET in response to Ca^2+^ elevations (**Fig. 3E**). These equilibrium shifts were observable for WT and both mutant PLB variants. Since we previously showed the rate of return of pentamer dissociation is rate-limiting for the return of PLB to SERCA after Ca^2+^ elevations, we compared the rate of PLB-SERCA rebinding for WT-, R9C-, and R14del-PLB. WT-PLB returned to SERCA with a τ of 3.5 ± 0.2 s, which was significantly slower than the rate of Ca^2+^ uptake (τ = 0.5 ± 0.03 s, *p* = 7.73E-8, **Fig. 3F**). R9C-PLB rebinding to SERCA was slower (τ = 4.8 ± 0.9 s), though the difference was not significantly different from WT (*p* = 0.28). These FRET changes are likely due to a population of non-crosslinked R9C-PLB that is still able to interact dynamically with SERCA with responsiveness to cytoplasmic Ca^2+^ changes. Interestingly, the rebinding of R14del-PLB to SERCA after Ca^2+^ elevations was significantly slower than WT-PLB (τ = 5.7 ± 0.8 s, *p* = 0.003) (**Fig. 3G, H**), suggesting that slow unbinding of R14del pentamers causes a considerable delay in the physiological exchange of PLB from its regulatory complex with SERCA during Ca^2+^ signaling. The kinetics of WT and mutant PLB rebinding SERCA after Ca^2+^ elevations are summarized in Supplementary **Fig. S6** and **Table S4**.

### Slow exchange of PLB monomers from R14del PLB pentamers disrupts Ca^2+^ regulatory dynamics

To understand the functional consequences of slower exchange of PLB pentamers containing DCM mutation, we modeled the responsiveness of SERCA regulation to changing Ca^2+^ over a physiological range of pacing frequencies (*19*). We considered that the enhanced stability of the PLB pentamer via a decrease in PLB-PLB K_D_ (**Fig. 1D,F**) could result from either an increase in the rate of pentamer oligomerization (*k*_*on*_) or a decrease in the rate of pentamer dissociation (*k*_*off*_), according to the relationship (*K*_*D*_ *= k*_*off*_*/k*_*on*_). Moreover, we observed that the exchange rate for the R14del mutation is slower (**Fig. 2**), and PLB pentamer exchange is the sum of the on- and off-rates, according to the relationship (*k*_*exchange*_ *= k*_*on*_*+k*_*off*_). Thus, we conclude that K_D_ is decreased as a result of a decrease in *k*_*off*_, rather than an increase in *k*_*on*_. Therefore, we simulated the effect of R14del mutation on PLB binding dynamics by slowing the unbinding rate of PLB pentamers by 2-fold to account for the observed 2-fold change in K_D_. **Fig. 4A** shows a simulation of WT- and R14del-PLB interaction with SERCA adjusting from a resting cardiac pacing frequency (60 bpm) to intense exercise (180 bpm) and then back to rest (60 bpm). At 60 bpm, the decreased dissociation rate of R14del pentamers reduced SERCA bound to PLB at resting heart rate (**Fig. 4B**) and increased the relative amount of PLB sequestered in pentamers (**Fig. 4C**). Thus, the computational model predicts that R14del causes SERCA to be dysregulated even at resting heart rates. Then, during exercise, SERCA becomes even more dysregulated as a larger fraction of PLB is sequestered into pentamers. Single-exponential decay fitting these changes in PLB-SERCA binding between rest and exercise demonstrated that the WT regulatory complex adjusts in and out of the exercising condition with a τ of ∼1s. However, with the pentamer off rate slowed 2-fold by R14del mutation, these adjustment times occur with a τ of ∼2s (**Fig. 4D,E**), suggesting that adjustments in PLB regulation between rest and exercise occur more slowly with R14del. Additionally, dynamic oscillations in PLB-SERCA binding during Ca^2+^ elevations were also decreased (**Fig. 4F**), indicating that the beat-to-beat changes in SERCA regulation were blunted by R14del mutation of PLB.

**Figure 4.**
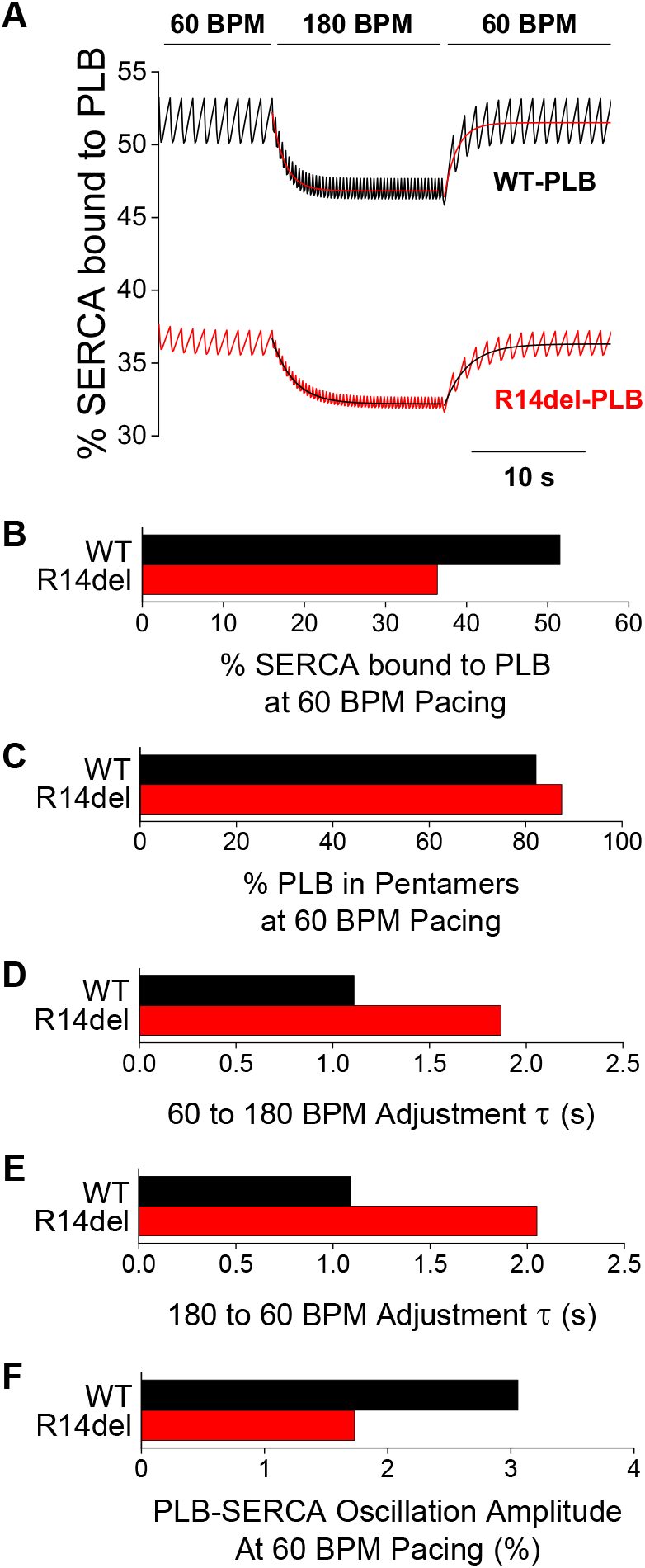
A computational model simulates PLB-SERCA complex exchange during changes in heart rate for WT and R14del PLB. **A)** A simulation of PLB-SERCA binding exchange during Ca^2+^ signaling adjusted from 60 bpm to 180 bpm to 60 bpm pacing rate for WT-(*black*) and R14del-PLB (*red*). Single-exponential decay fits reveal characteristic time constants (τ) for adjustment of PLB-SERCA binding equilibria between 60 and 180 bpm pacing frequencies. **B)** Quantification of the percent SERCA bound to PLB at 60 bpm. **C)** Quantification of the percent PLB in pentamers at 60 bpm **D)** Quantification of the adjustment τ of PLB-SERCA from 60 to 180 bpm. **E)** Quantification of the adjustment τ of PLB-SERCA from 180 to 60 bpm. **F)** Quantification of the oscillation amplitude of PLB-SERCA complexes unbinding and rebinding in response to Ca^2+^ elevations during 60 bpm pacing.

## DISCUSSION

### R14del mutation stabilizes PLB pentamers

The naturally occurring R14del mutation of PLB was originally discovered in a large family in which carriers experienced lethal hereditary dilated cardiomyopathy by middle age. The seminal first study of this mutation showed that coexpression of R14del-PLB resulted in multiple PLB pentamer bands with different mobilities on Western blot. This was initially taken to indicate destabilization of the PLB pentamer by co-expression of R14del-PLB (*5*). That conclusion contrasts with the model proposed here, that R14del-PLB *increases* the stability of the PLB pentamer. We also observed multiple bands for PLB pentamers by Western blot (**Fig. 1A**), but these bands are present for both WT-PLB and R14del-PLB oligomers. This pattern of mobility may arise from differential degrees of phosphorylation of 0 to 5 PLB protomers in the pentamers (*16, 33*). Moreover, R14del mutation resulted in an apparent increase in the ratio of pentamers to monomers (**Fig. 1A**), an increased pentamer melting temperature (**Fig. 1B**), and a higher affinity for homo-oligomerization in live cells (**Fig. 1D, F**), consistent with the interpretation that the R14del mutation increases PLB pentamer stability.

The expected functional consequence of increased PLB pentamer stability is a decrease in the population of PLB monomers available to inhibit SERCA. This prediction comports with the present observation of decreased binding of R14del-PLB to SERCA (**Fig. 1G,H**). Although early reports suggested a gain-of-function character for R14del-PLB, resulting in superinhibition of SERCA (*5*), several subsequent biochemical studies found that R14del-PLB expressed alone (*20, 21, 23, 29*) or in combination with WT (*21, 23-25*) led to reduced SERCA inhibition compared to WT-PLB. The results of the present study may provide a possible mechanism for the observed loss-of-function character of the R14del mutation, and the dominant negative effect of R14del when coexpressed with WT-PLB (**Fig. 1**), since we observed increased oligomerization of R14del-PLB alone and increased oligomerization of R14del-PLB with WT-PLB (**Fig. 1D,F**). By sequestering both mutant and WT-PLB away from SERCA, R14del-PLB has a dominant negative effect that suppresses PLB inhibition of Ca^2+^ transport in heterozygous carriers. This characteristic of the R14del mutation is reminiscent of another DCM mutation of PLB, R9C, which also co-oligomerizes with WT-PLB in stable hetero-pentamers (*22*). We speculate that oligomerization of micropeptides starts with formation of dimers, then additional protomers are recruited onto that template to form the rest of the higher order oligomers (*34*). Thus, mutations that increase the stability of the initial dimer have a larger than expected effect in assembling mixed mutant/WT oligomers. In contrast to consistent observations in biochemical studies of R14del-PLB, the functional consequence of R14del-PLB expression *in vivo* has been less clear. Some physiological studies of R14del-PLB-expressing mice reported weak Ca^2+^ handling and decreased cardiac contractility (*35-37*) (suggesting R14del-PLB is superinhibitory). In contrast, other studies showed increased SERCA activity, with larger Ca^2+^ transients, enhanced cardiac contractility, and faster relaxation (*25, 38*) (suggesting R14del-PLB is less inhibitory than WT-PLB). A similar disparity exists in biochemical and physiological studies of the R9C mutation of PLB, which showed a loss-of-inhibition character *in vitro* (*6, 12, 20-22, 39*) but caused diminished calcium handling *in vivo* (*6, 39*). One explanation for these disparate reports is that there may be a difference between acute effects detected in *in vitro* experiments versus chronic downstream consequences at later stages *in vivo*. Acute studies may illuminate the fundamental pathological mechanism while long term expression may be more representative of the human disease state after full evolution of pathological mechanisms, compensatory physiological adaptation, and cardiac remodeling (*11, 12*).

### R14del PLB pentamers exchange slowly and are less responsive to changing physiological demands

The rate of Ca^2+^ transport by SERCA is a key determinant of the strength of cardiac contraction (inotropy) and the speed of cardiac relaxation (lusitropy). Dynamic adjustment of Ca^2+^ handling enables the heart to increase cardiac output to meet the stress of exercise and then decrease physiological performance after returning to rest. Some of this dynamic modulation of SERCA function is achieved through adrenergic signaling, as discussed below, but there are also important intrinsic mechanisms that contribute to the response. In particular, an increase in heart rate results in an increase in contraction strength. This phenomenon is known as treppe, positive staircase, or the Bowditch effect (*19, 40-43*). Loss of this positive force-frequency relationship is a hallmark of heart failure in patients (*43, 44*). We have previously proposed that the dynamic exchange of PLB between SERCA-bound and pentamer-bound pools contributes to the Bowditch effect. Specifically, we found that a fraction of the population of PLB bound to SERCA is liberated during Ca^2+^ elevations and joins the pentameric pool of PLB, then it unbinds from pentamers and rebinds SERCA when Ca^2+^ concentration returns to baseline (*19, 45*). Because of the high temporal stability of PLB pentamers (*18, 19*) (**Fig. 2**), more frequent Ca^2+^ elevations (at higher heart rates) cause accumulation of PLB in the pentameric form, resulting in less inhibition of SERCA relative to slow heart rates. Thus, the degree of sequestration of PLB in non-inhibitory pentamers is dependent on cardiac pacing frequency (heart rate) (*19*). Here we find that R14del-PLB pentamers are even more stable than WT-PLB pentamers, causing protomers to unbind more slowly after cellular Ca^2+^ elevations (**Fig. 2**). A computational model suggested that this resulted in an impaired Bowditch mechanism. The R14del mutation caused the PLB-SERCA regulatory complex to respond sluggishly to changing heart rates (**Fig. 4D-E**) and it also blunted changes in PLB-SERCA binding on a beat-to-beat basis (**Fig. 4F**). The reduced responsiveness of R14del pentamers to these physiological cues may explain the decreased responsivity of Ca^2+^ handling to changes in pacing frequency observed in cardiomyocytes with heterozygously expressing R14del-PLB (*24, 25*). This pathological mechanism may also be operative for the R9C mutation, but we were unable to detect any exchange of R9C-PLB pentamers during cellular Ca^2+^ signals, probably because oxidative cross-linking of R9C-PLB (*22*) largely prevented exchange of protomers (**Fig. 5**). Thus, R9C-PLB is even less responsive than the R14del mutant and may contribute to the more severe phenotype of R9C. The average survival for carriers of R9C is estimated at 25 years (*6*) versus the estimated 38 years for carriers of R14del (*9*).

**Figure 5.**
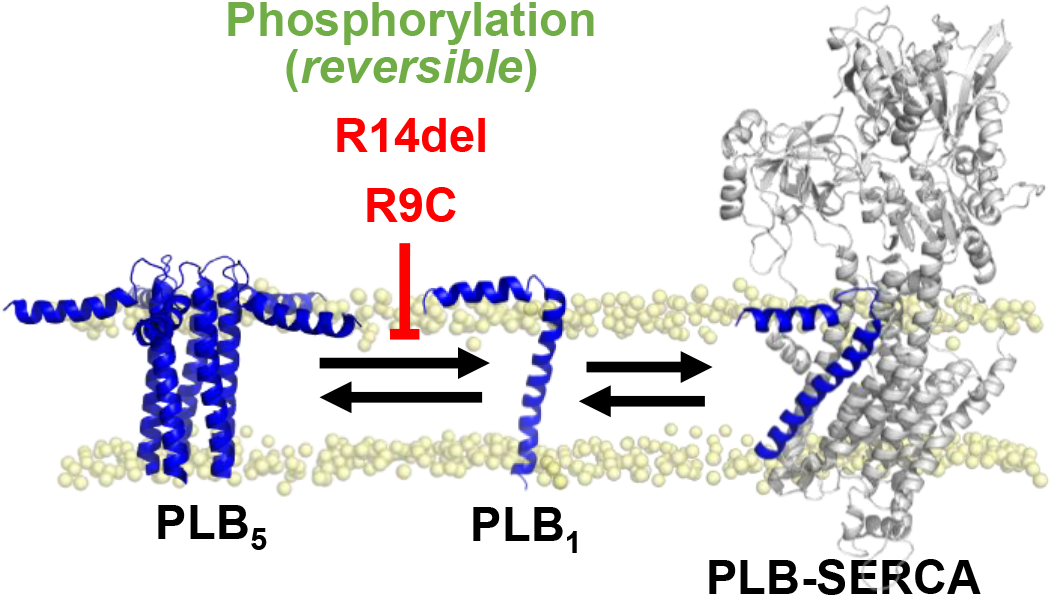
A schematic diagram displaying how PLB phosphorylation and DCM mutations of PLB, R14del and R9C, impact the equilibrium binding of monomeric PLB (PLB_1_) with pentamers (PLB_5_) and SERCA (PLB-SERCA) by decreasing the rate of PLB monomers unbinding from pentamers.

### Impaired PLB binding dynamics may contribute to arrhythmogenesis

In addition to DCM, R14del-PLB is also associated with another pathological condition, arrhythmogenic right ventricular cardiomyopathy (ARVC). Human carriers of R14del mutation experience arrhythmias that often begin before the onset of heart failure and can lead to sudden cardiac death (*46, 47*). Although ARVC is more commonly associated with mutations in proteins of the intercalated disc (*48, 49*), it has been proposed that R14del-PLB and these genetic perturbations of intracellular integrity may lead to the development of arrhythmia through a common pathogenic mechanism. Specifically, disruptions of the desmosome in ARVC have been found to result in a maladaptive changes in Ca^2+^ handling by increasing PLB phosphorylation (*49, 50*). While phosphorylation normally causes a temporary increase in SERCA activity during physiological stress that is readily reversible during rest, chronic hyperphosphorylation of PLB after desmosome disruption may lead to SR Ca^2+^ overload and premature ventricular contractions (*50*). Indeed, increased PLB phosphorylation has been reported across various models of arrhythmogenic cardiomyopathy (44-46). Interestingly, like R14del mutation, PLB phosphorylation also increases the oligomerization (16, 17) and temporal stability of pentamers (19). Thus, we speculate that enhanced pentamer stability and impaired PLB binding dynamics, caused by either hyperphosphorylation of PLB or R14del mutation, may serve as common arrhythmogenic mechanisms in ARVC.

### Summary

**Fig. 5** illustrates the principal finding of this study, that the R14del mutation of PLB causes hyper-stabilization of PLB pentamers and blunts the dynamics of regulation of cardiac calcium handling by PLB. This is similar to the reversible stabilization of pentamers induced when PLB is phosphorylated or the covalent crosslinking of PLB oligomers that occurs with R9C mutation of PLB. Decreasing the off-rate of PLB monomers from pentamers depletes the overall level of free monomers, decreasing the fraction of SERCA regulated by PLB. In addition, excessive stabilization of the oligomer also slows the release of PLB protomers into the active monomeric pool as cytosolic calcium levels fall during diastole. This results in sluggish transitions between the different regulatory stances of exercise versus rest, and it damps the beat-to-beat changes in regulation of SERCA by PLB. We propose that this poor responsiveness to physiological stress stimulates cardiac remodeling and eventual failure of the heart in dilated cardiomyopathy.

## METHODS

### Cell culture and polyethylenimine-mediated (PEI) transfection

HEK-293 were obtained from American Type Culture Collection and were maintained in an incubator (37°C, 5% CO_2_) in Dulbecco’s modified eagle medium (DMEM, Cat#219-010-XK, Wisent Bioproducts, Saint-Jean-Baptiste, QC, Canada), supplemented with 10% fetal bovine serum (ThermoFisher Scientific, Cat#12483020), penicillin-streptomycin (Cat:10378016, 1:100 dilution), and non-essential amino acid solution (Cat#10378016, 1:100 dilution). For transfection, cells were split, so that they reached 70∼80% confluency on the day of transfection. Transfection reagents were prepared by mixing 1 mL serum-free DMEM, 5 μg of plasmid DNA, and 15 μg PEI (Polysciences Inc., Cat#23966-1, Warrington, PA) in a 1.5 mL Eppendorf tube, vortexed and rest at the room temperature for 20 minutes. Transfection reagents were added dropwise to cells and transfected cells were returned to an incubator overnight. Next day, 10 mL fresh DMEM + 10% FBS were given to cells and incubated for another day.

### Immunoblotting and protein thermostability assays

Following transfection, cells were washed twice with ice-cold 1x phosphate-buffered saline (pH7.4; 137 mM NaCl, 2.7 mM KCl, 8 mM Na_2_HPO_4_, 2 mM KH_2_PO_4_) and protein lysates were harvested in 8 M urea (BioShop, URE001, Toronto, Canada), supplemented with protease inhibitors (Roche, 04574834001). Protein lysates were sonicated. Insoluble fractions were spanned down at 15,000x*g* for 15 minutes at room temperature and cell-free lysates were transferred to a new Eppendorf tube. Protein concentration was qualified by Bradford assays (Sigma-Aldrich, B6916-500 mL). 30 μg soluble proteins were resolved on a 15% sodium dodecyl sulphate poly acrylamide gel via electrophoresis (SDS-PAGE) at 90 V, followed by electrophoretic transfer of proteins to a 0.22 μm nitrocellulose membrane (Bio-Rad, Cat#1620112) at 65 V for 60 minutes. A membrane was blocked with 5% non-fat milk in Tris-buffer saline (TBST, 20 mM Tris-HCl; pH7.4, 150 mM NaCl, 0.1% (w/v) Tween 20). 0.1 μg/mL anti-PLB antibody (ThermoFisher, Cat#MA3-922) in TBST was added to the membrane and incubated overnight at 4°C on a rotor. Next morning, the membrane was washed 3 time with TBST. HRP-conjugated anti-mouse antibodies (Promega, Cat#W4021) in TBST was added to membranes and rocked on a rotator for 1 hour at the room temperature. Membranes were washed 3 time with TBST, incubated with chemiluminescent reagents (ThermoFisher, Cat# 32209) in the dark for 5 minutes. Immunoblot signals were captured by ChemiDoc (Bio-Rad). Densitometry analysis of signal intensity was carried out in Image Lab software (Bio-Rad) as per manufacturer’s instruction.

For protein thermostability assays, cell-free lysates in 8 M urea were thoroughly mixed with protein loading dye before heated up in a range of temperature (25∼70°C, 5°C increments) for 2 minutes in a thermocycler. Lysates were then subjected to SDS-PAGE runs. % PLB oligomers was estimated as the following. Percentage PLB oligomers = Avg. oligomer density / (Avg. monomer density + Avg. oligomer density) * 100%.

### FRET acceptor-sensitization in HEK-293 cells

Acceptor sensitization FRET was quantified as previously described (*22*). Briefly, HEK-293 cells were transiently transfected with mCer-donor and YFP acceptor–labeled FRET binding partners in a 1:5 M plasmid ratio. Transfected cells were reseeded to a poly-D-lysine coated chamber slide and imaged using an inverted microscope (Nikon Eclipse Ti2). Field of view images of cells were acquired 24-48 hours post transfection with a 20X objective, numerical aperture 0.75. A Lumencor Spectra X excitation system was used to excite samples with 50 ms exposure time for mCer donor (excitation: 420/440 nm, detection: 475 nm), YFP acceptor (excitation: 510/525 nm, detection: 540 nm), and FRET (excitation: 420/440 nm, detection: 540 nm) channels. Emitted light was passed through a dichroic emission filter cube (CFP/YFP/mCherry Spectra X emission filter set) before detection with a Photometrics Prime 95B 25 mm camera. Seventy-two images were collected for each channel in a 9 × 9 grid with a 1600 μm step size between each image. For each condition, two sets of images (yielding ∼500 total cells per condition) were collected from 5 independent experiments. Whole cell fluorescence intensity was quantified from images using automated analysis with a custom script in FIJI software. Specifically, cells with a minimum mCer intensity of 250 AU and an area of 136 – 679 μm2 with at least 40% circularity were selected for high throughput cell scoring with a rolling background subtraction. Fluorescence intensities from mCer, YFP, and FRET channels (*I*_*DD*_, *I*_*AA*_, and *I*_*DA*_ respectively) were used to calculate sensitized emission FRET according to the formula *E*_*app*_ *= F*_*c*_ */ (F*_*c*_ *+ G x I*_*DD*_), where *F*_*c*_ *= I*_*DA*_ *- (a x I*_*AA*_*) – (d x I*_*DD*_). Here, *F*_*c*_ represents the sensitized emission FRET intensity corrected for crosstalk between channels and *E*_*app*_ represents the apparent FRET efficiency corrected for imaging induced photobleaching. The parameters *a* and *d* are crosstalk constants calculated as *a = I*_*DA*_*/I*_*DD*_ for a control sample transfected only with the YFP acceptor and *d = I*_*DA*_*/I*_*AA*_ for a control sample transfected only with the mCer donor. *G* is the ratio of sensitized acceptor emission to a corresponding amount of donor recovery in the *I*_*DD*_ channel after acceptor photobleaching (*I*_*DD*_^*post*^), defined by the equation *G = F*_*c*_*/(I*_*DD*_^*post*^*-I*_*DD*_) (*51*). For the experiments in this study using mCer and YFP FRET pairs, these values were determined to be *a* = 0.1853, *d* = 0.4051, and *G* = 2.78. Apparent FRET efficiencies for each cell were then plotted as a function of YFP acceptor fluorescence intensity, which was used as an index of relative protein expression (*22*). This plot yielded a FRET-based binding curve, illustrating the relative concentration dependence of *E*_*app*_. FRET efficiency was low in cells with low protein expression, increasing to a maximum in the cells with the highest protein expression. The data were fit with a hyperbolic function of the form *FRET = FRET*_*max*_ *x Protein / (K*_*D*_ *+ Protein)*, where *FRET*_*max*_ is the maximal FRET efficiency at high protein concentration (representing the intrinsic FRET efficiency of the bound complex), *Protein* is inferred from the relative YFP fluorescence intensity in each cell, and *K*_*D*_ is the relative dissociation constant (the protein concentration that yields half-maximal FRET efficiency). The *K*_*D*_ is inversely related to the relative affinity of the protein-protein interaction. For PLB-PLB binding curves, the FRET_max_ value for each condition was shared among data from 5 independent experiments, giving 5 independent K_D_ values for each group. Differences in the apparent K_D_ values between groups were determined by one-way ANOVA with Tukey’s *post-hoc* test (*= *p*>0.05). Differences in the FRET_max_ and K_D_ values for SERCA-PLB binding for WT-PLB and R14del-PLB were determined by student’s t-test (*= *p*>0.05).

### Confocal fluorescence microscopy to measure intermolecular FRET and intracellular Ca^2+^

HEK-293 cells exhibiting spontaneous Ca^2+^ oscillations were generated by transient transfection with GFP-RyR2 and either Cer or unlabeled SERCA2a and co-transfected with WT or mutant PLB FRET pairs tagged with Cer and YFP fluorescent proteins. Transfected cells were cultured for 24 h and seeded into poly-D-lysine coated glass bottom chamber slides in DMEM plus 10% fetal bovine serum. Twenty-four hours after seeding, cell culture medium was changed with PBS (+Ca^2+^/+Mg^2+^), and experiments were conducted with a Zeiss LSM 880 confocal microscope using a 40× oil immersion objective. To observe transient changes in cytoplasmic calcium, cells were incubated with 2 μM X-Rhod-1/AM (X-Rhod) for 20 min in PBS (+Ca^2+^/+Mg^2+^). X-Rhod was excited with the 594 nm line of a He-Ne laser, and emitted fluorescence was measured at wavelength 580 nm. FRET pair fluorophores Cer and YFP were excited with the 458 nm line of an argon laser, and emitted fluorescence was measured at wavelengths 485 ± 15 and 537 ± 15 nm, respectively. Images were acquired in line scan every 24 ms for ∼2 min. FRET ratio was determined by dividing the acceptor fluorescence by the donor fluorescence and plotted as a function of time with X-Rhod to indicate concurrent changes in [Ca^2+^]. FRET ratio data was smoothed using a Savitzky–Golay binomial filter with a 4.08 s averaging window. Changes in FRET ratio and X-Rhod fluorescence associated with Ca^2+^ uptake were fit to the single-exponential decay function, γ = A1^−x/τ^ + γ_0_ in Origin, to estimate the time constant (τ) of the change, where A1 is the amplitude of change and γ_0_ is the initial detected fluorescence. Differences in the τ between groups were determined by one-way ANOVA with Dunn’s Sidak *post-hoc* test (*= *p*>0.05).

### Generation of a model of the pentameric form of R14Del PLB

A structural model of the mutant pentamer was created using the NMR structure of the WT-PLB pentamer (PDB: 2KYV) (*13*). Among the 20 states included in this structure, we selected the three states that present the least number of Phi and Psi torsion angles in disfavored Ramachandran’s plot areas, namely states 04, 16 and 17. Starting from these structures, we built three R14del models using VMD’s molecular editing plugin Molefacture (*52*). R14 atoms were removed, and residues 1-13 were attached to residues 15-52. To preserve the helical structure of the cytoplasmic domain in R14del-PLB models, we transferred the omega angle between WT R14-C and A15-N atoms to the peptide bond between R14del R13-C and A15-N atoms. The six resulting models (three WT PLB structures and three R14del structures) were incorporated into POPC membranes and two 10Å thick slabs of 0.15 M [KCl] solution, using the webserver CHARMM-GUI (*53*). The Molecular Dynamics (MD) simulations were conducted in two phases, using NAMD program (*54*) and CHARMM36m forcefield (*55*). A switch potential was used to smoothly reduce 8Å-12Å long-range non-bonded energies to 0. To comply with the applied periodic boundary conditions, long-range electrostatic energies were calculated using Particle Mesh Ewald algorithm. The first phase of equilibration was divided into four steps, each in NPT conditions. The first step was 1ns long, using a timestep of 1 fs. It involved specific harmonic constraints applied on all PLB atoms and on POPC phosphorus atoms (2 kcal.mol^-1^). The following steps were simulated using a 2 fs timestep. During the 13ns long second step, the harmonic constraints applied on PLB backbone atoms during step 1 were maintained, while those applied on PLB sidechain atoms and POPC were reduced to 1 kcal.mol^-1^. During the 20ns long third step, only the harmonic those applied on PLB backbone atoms were maintained at 1 kcal.mol^-1^. The last fourth step was performed without any constraints for 20ns. The production phase was conducted in NPT conditions, using a Langevin thermostat to maintain the temperature of the MD system at 300K coupled with a Langevin Piston to keep the pressure fixed at 1 bar, for a duration of 1μs. The SHAKE algorithm (*56*) was used to fix the bond lengths that involve hydrogens.

MD analyses consisted in monitoring the PLB backbone atoms’ RMSD, PLB Cα atoms’ RMSF, and the displacement of the tilt angle between the axis of PLB transmembrane bundle and the axis normal to the membrane every 2 ns throughout each production MD trajectory. The calculations were conducted with TCL written programs. To facilitate their interpretation, results were averaged over the entire WT and R14del datasets, respectively, using R written scripts. Time-lag dependent Pearson autocorrelation coefficients were calculated for the time-dependent tilt angles values using the ACF function (*57*) included in the ‘stats’ R package. To facilitate the interpretation of the autocorrelation profiles, the obtained autocorrelation profiles were fitted with exponential decay functions. Solvent-accessible electric potentials were calculated using APBS online server (*58*). The model used are the centroid models of the most populated K-means group among WT-PLB and R14del-PLB datasets, respectively. K-means clustering was conducted using CPPTRAJ program included in Amber18 package (*59*).

### Kinetic modeling

We utilized a computational model that we previously developed to describe the exchange of WT-PLB monomers between SERCA- and pentamer-bound pools in response to Ca^2+^ signaling (*19*). Briefly, a series of ordinary differential equations was used to describe the binding and unbinding kinetics of these membrane protein complexes under diastolic and systolic conditions. The mean rate constants for WT-PLB were previously determined or constrained based on fitting Ca^2+^-dependent population shifts in PLB-SERCA binding measured by FRET using a genetic algorithm (*19, 60*). For simulations of R14del-PLB, the off-rate of the PLB pentamer (PLB unbinding from pentamers) from the original model was decreased two-fold to reflect the two-fold decrease in PLB-PLB K_D_ with R14del-mutation observed from FRET-based binding curves (**Fig. 1F**) which resulted in a slower rate of pentamer unbinding (**Fig. 2D**). For simulations of PLB binding dynamics during cardiac pacing, simulated Ca^2+^ transients were used as inputs with varying pacing frequency. The ordinary differential equation system was numerically solved using the scipy function (v1.5.0) SOLV_IVP. Adjustments in the population of PLB-SERCA were well described by a single-exponential decay function, γ = A1^−x/τ^ + γ_0_ in OriginLabs software, to estimate the time constant (τ) of the change, where A1 is the amplitude of change and γ_0_ is the initial value of this population in the model.

## ACKNOWLEDGEMENTS

The authors would like to acknowledge Ava Vandenbelt for technical assistance. The authors would also like to thank Antonio Zaza and Francesco Lodola for helpful discussions. This investigation was supported by the Heart and Stroke Foundation of Canada: G-17-0016337 to A.O.G. This study was also supported by the National Institutes of Health (NIH): Maximizing Investigators’ Research Awards (MIRA) R35GM124977 from the National Institute of General Medical Sciences (NIGMS) to P. M. K.-H.; Ruth L. Kirschstein Predoctoral Individual National Research Service Award (NRSA) F31HL165900-01 from the National Heart, Lung, and Blood Institute (NHLBI) to S.R.C; R01HL092321 and R01HL143816 from the NHLBI to S. L. R.

