## Supplementary material for "Dilated cardiomyopathy variant R14del increases phospholamban pentamer stability, blunting dynamic regulation of cardiac calcium handling": Table S1

**Supporting Information**

| PLB-PLB $K_D$ analyzed by 1-way ANOVA with Tukey's post-hoc | WT-WT | R9C-R9C | WT-R9C | R14del-R14del |
|---|---|---|---|---|
| WT-R14del | **$9.85 \times 10^{-7}$*** | **$2.50 \times 10^{-3}$*** | **$3.27 \times 10^{-5}$*** | **$8.51 \times 10^{-4}$*** |
| R14del-R14del | **$1.48 \times 10^{-8}$*** | **$1.30 \times 10^{-7}$*** | **$2.66 \times 10^{-8}$*** | |
| WT-R9C | 0.44 | 0.32 | | |
| R9C-R9C | **0.01*** | | | |

**Supplementary Table S1.** *P* values comparing the differential dissociation constants ($K_D$s) of PLB-PLB binding for wild-type- and mutant- PLB pentamer oligomerization. For each pair, the first binding partner is the FRET-donor, and the second is the acceptor. Data were analyzed by 1-way ANOVA with Tukey's *post-hoc* test ($p<0.05$ = *****).
