## Supplementary material for "Dilated cardiomyopathy variant R14del increases phospholamban pentamer stability, blunting dynamic regulation of cardiac calcium handling": Fig. S1

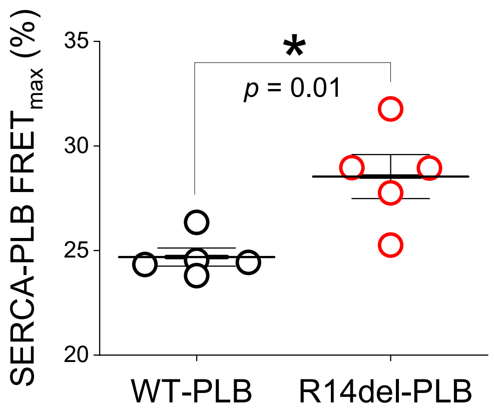


**Supplementary Figure S1.** $FRET_{max}$ values for SERCA- WT-PLB and R14del-PLB determined from FRET-based binding curves. Differences were determined by student's t-test ($p<0.05$ = *).
