## Supplementary material for "Dilated cardiomyopathy variant R14del increases phospholamban pentamer stability, blunting dynamic regulation of cardiac calcium handling": Fig. S2

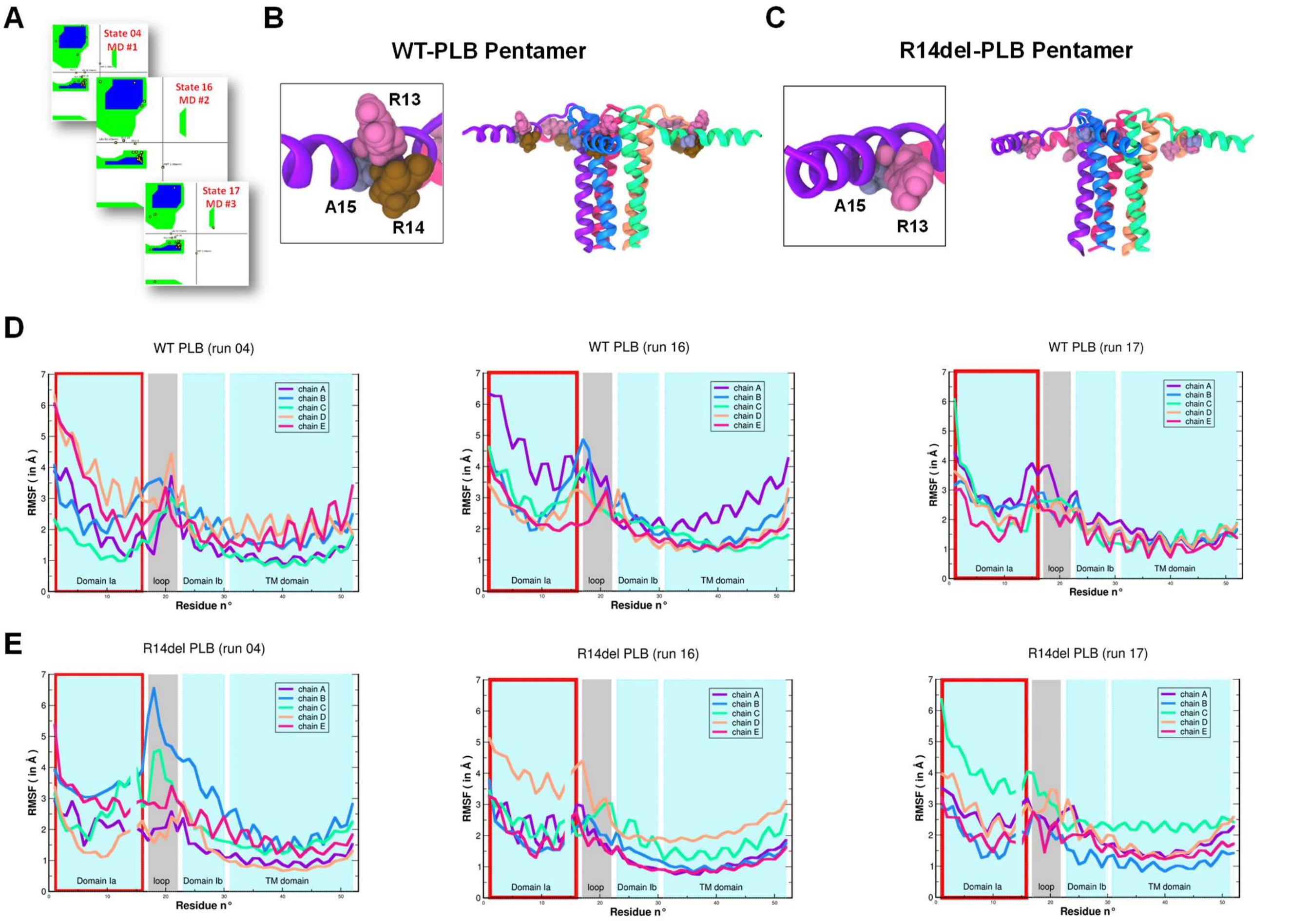


**Supplementary Figure S2.** Molecular dynamics simulations of WT- and R14del-PLB pentamer structures. **A)** Three distinct NMR structures (state 04, state 16, and state 17) with the least number of incorrect torsion angles on Ramachandran plots were selected from 20 published NMR structures of the WT-PLB pentamer (PDB:2KYV) for MD simulations. **B,C)** Models of the R14del pentamer (panel C) were generated by removing Arg14 from the WT structure (panel B, *brown*) and bridging the ends of Arg13 (*pink*) and Ala15 (*grey*) while maintaining the value of the torsion angle omega for the 14-15 peptide bond. **D)** Evolution of the WT-PLB backbone flexibility as measured by RMSF. Colors for individual PLB subunit chains correspond to the colors of subunits shown in panel B. **E)** Evolution of the R14del-PLB backbone flexibility as measured by RMSF. Colors for individual PLB subunit chains correspond to the colors of subunits shown in panel C.
