## Supplementary material for "Dilated cardiomyopathy variant R14del increases phospholamban pentamer stability, blunting dynamic regulation of cardiac calcium handling": Fig. S3

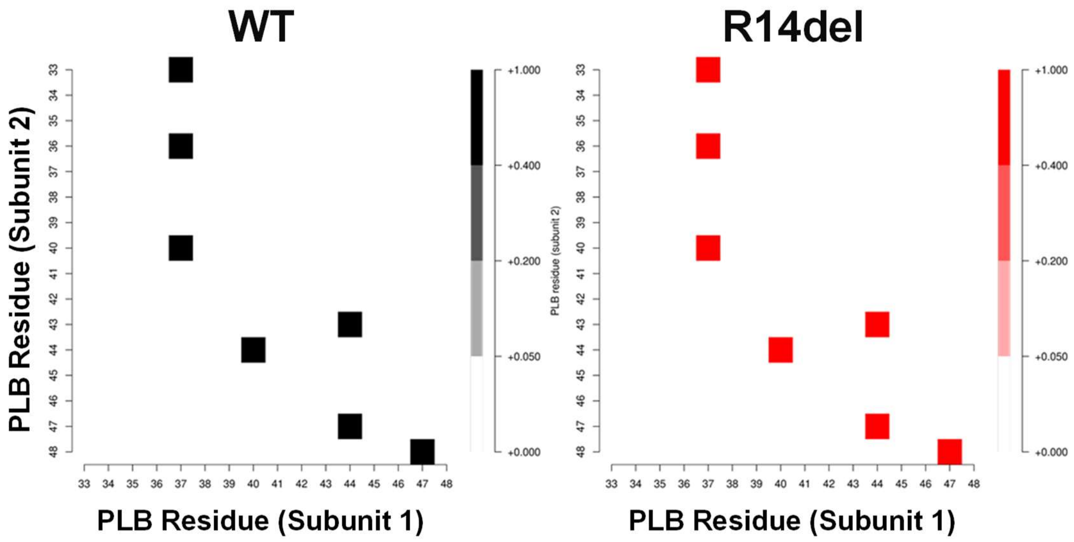


**Supplementary Figure S3.** Contact map of PLB TM residue inter subunit interactions for WT (left, *black*) and R14del (right, *red*) -PLB.
