## Supplementary material for "Dilated cardiomyopathy variant R14del increases phospholamban pentamer stability, blunting dynamic regulation of cardiac calcium handling": Fig. S4

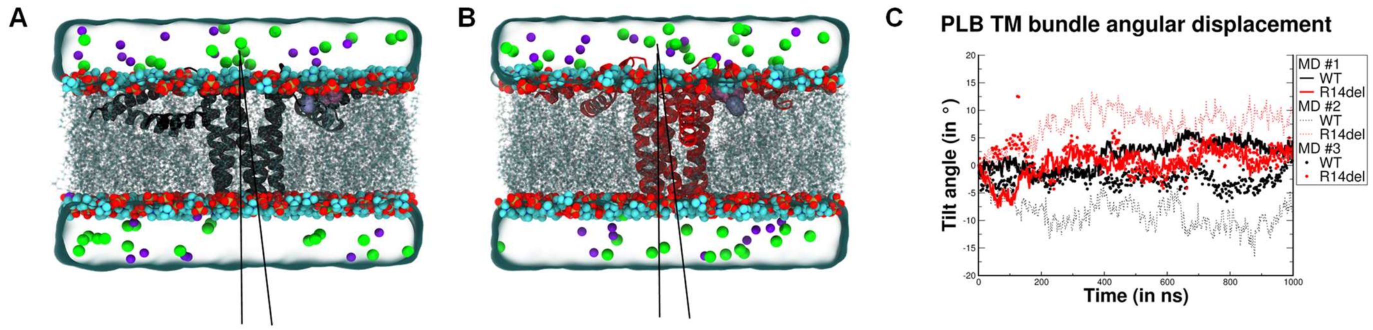


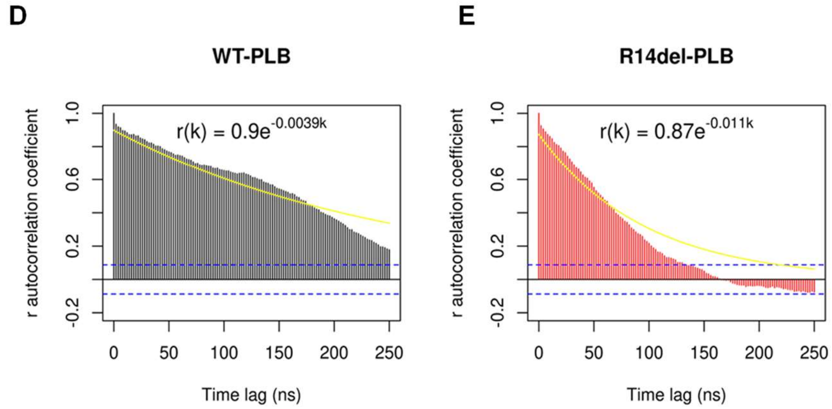


F

| MD system | Exponential decay rates | |
|---|---|---|
| | WT | R14del |
| MD #1 | -0.0039 * | -0.011 * |
| MD #2 | -0.015 | -0.008 |
| MD #3 | -0.023 | -0.027 |
| Average | -0.014 +- 0.01 | -0.015 +- 0.01 |

**Supplementary Figure S4.** Molecular Dynamics analyses of the transmembrane (TM) domain bundle of PLB pentamers. **A)** Simulation box of the most populated cluster representative of WT-PLB. **B)** Simulation box of the most populated cluster representative of R14del-PLB. In panels A and B the tilt angle of the TM bundle is depicted as a gray triangle. **C)** Time-dependent tilt angle displacement of WT-PDB and R14del-PLB pentamers MD trajectories #1, #2, and #3 were generated from MD simulations of PDB: 2KYV states 04, 16 and 17, respectively. **D,E)** The time-lag dependent Pearson "r" autocorrelation coefficients of WT-PLB and R14del-PLB angle displacements, respectively, throughout MD #1. On both panels, the exponential decay function curve is depicted as a yellow line, and the equation function is displayed in the top side of the plot. **F)** Summary of the exponential decay rates calculated for the time-lag dependent TM bundle tilt angle autocorrelation in MD simulations #1 (highlighted in yellow), #2, and #3 and the average decay rates for WT and R14del PLB.
