## Supplementary material for "Dilated cardiomyopathy variant R14del increases phospholamban pentamer stability, blunting dynamic regulation of cardiac calcium handling": Fig. S5

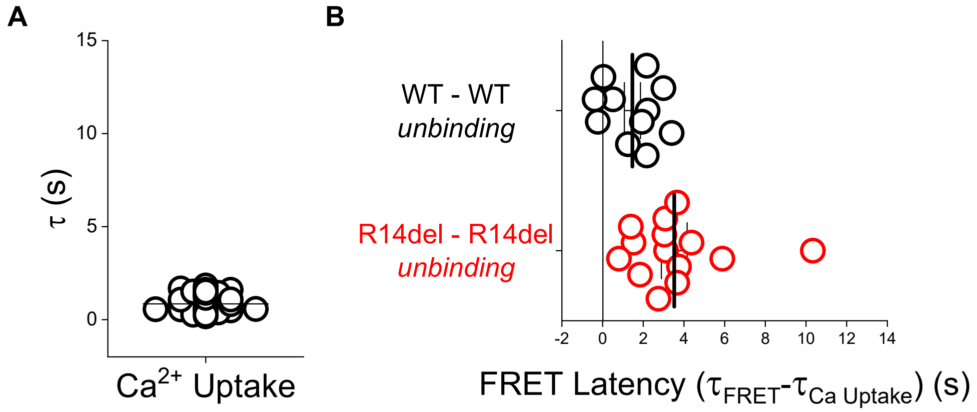


**Supplementary Figure S5. A)** Time constants for $Ca^{2+}$ Uptake quantified for experiments measuring PLB-PLB binding dynamics in response to $Ca^{2+}$ signaling. **B)** The latency of PLB-PLB FRET ratio changes compared to $Ca^{2+}$ Uptake ($\tau_{FRET} - \tau_{Ca\ Uptake}$) with lines representing mean ± SEM.
