## Supplementary material for "Dilated cardiomyopathy variant R14del increases phospholamban pentamer stability, blunting dynamic regulation of cardiac calcium handling": Table S2

| Apparent τ (Mean ± SEM) | |
|---|---|
| **Process** | **τ (s)** |
| WT – WT (*unbinding*) | 2.6 ± 0.4 |
| R14del – R14del (*unbinding*) | 4.4 ± 0.7 |
| $Ca^{2+}$ Uptake | 1.0 ± 0.1 |

**Supplementary Table S2.** Time constants (τ) quantified for the rate of WT- and R14del- PLB-PLB unbinding associated with $Ca^{2+}$ uptake. Time constant values are reported as mean ± SEM.
