## Supplementary material for "Dilated cardiomyopathy variant R14del increases phospholamban pentamer stability, blunting dynamic regulation of cardiac calcium handling": Table S3

| PLB-PLB unbinding analyzed by 1-way ANOVA with Dunn's post-hoc | | |
|---|---|---|
| | WT-WT (*unbinding*) | R14del-R14del (*unbinding*) |
| $Ca^{2+}$ Uptake | **0.01*** | **$4.87 \times 10^{-8}$*** |
| R14del-R14del (*unbinding*) | **0.01*** | |

**Supplementary Table S3.** *P* values comparing the differences in the time constant (τ)s for of PLB oligomer unbinding for WT- and R14del-PLB pentamers in response to $Ca^{2+}$ uptake. Data were analyzed by 1-way ANOVA with Dunn's *post-hoc* test ($p<0.05$ = *).
