## Supplementary material for "Dilated cardiomyopathy variant R14del increases phospholamban pentamer stability, blunting dynamic regulation of cardiac calcium handling": Table S4

| Apparent τ (Mean ± SEM) | |
|---|---|
| **Process** | **τ (s)** |
| WT – SERCA (*rebinding)* | 3.5 ± 0.2 |
| R9C – SERCA *(rebinding)* | 4.8 ± 0.9 |
| R14del – SERCA *(rebinding)* | 5.7 ± 0.8 |
| $Ca^{2+}$ Uptake | 0.5 ± 0.03 |

**Supplementary Table S4.** Time constants (τ) quantified for the rate of WT- and R14del- rebinding associated with $Ca^{2+}$ uptake. Time constant values are reported as mean ± SEM.
