## Supplementary material for "Dilated cardiomyopathy variant R14del increases phospholamban pentamer stability, blunting dynamic regulation of cardiac calcium handling": Table S5

| PLB-SERCA rebinding analyzed by 1-way ANOVA with Dunn's post-hoc | WT-SERCA (*rebinding*) | R9C-SERCA (*rebinding*) | R14del-SERCA (*rebinding*) |
|---|---|---|---|
| $Ca^{2+}$ Uptake | **7.73 x 10$^{-8}$*** | **1.02 x 10$^{-10}$*** | **2.67 x 10$^{-15}$*** |
| R14del-SERCA (*rebinding*) | **0.003*** | 0.63 | |
| R9C-SERCA (*rebinding*) | 0.28 | | |

**Supplementary Table S5.** *P* values comparing the differences in the time constant (τ)s for of PLB-SERCA rebinding for WT-, R9C-, and R14del-PLB in response to $Ca^{2+}$ uptake. Data were analyzed by 1-way ANOVA with Dunn's *post-hoc* test ($p<0.05$ = *****).
